# Tritrophic Interactions among Crapemyrtle, Crapemyrtle Aphids, and the Green Lacewing C*hrysoperla rufilabris* (Burmeister)

**DOI:** 10.64898/2026.11.24.766821

**Authors:** John J. Herbert, Romain Exilien, Heather McAuslane, R. F. Mizell

## Abstract

Tritrophic interactions can have considerable impact on biological control systems, therefore, it is imperative to consider these interactions when implementing or evaluating biological control agents. Crape myrtles, *Lagerstroemia indica* L. x *L. fauriei* L., are important non-native ornamental plants from Asia that are widely planted and a major woody landscape species in the southern U.S. Six alkaloids have been identified from *L. indica,* mostly in the seeds and leaves. Crapemyrtle aphids, *Tinocallis kahawaluokalani* (Kirkaldy), are host specific to crape myrtle and have been shown to be important prey for many native beneficial insects including coccinellids, mirids, and green lacewings. The crapemyrtle aphids occur in different densities per leaf on crape myrtle cultivars and some native beneficial insects cannot fully develop by feeding solely on these aphids. Moreover, no parasitoids of crapemyrtle aphids are known. This information suggests that plant chemistry is involved in tritrophic interactions between this plant, its herbivores and their natural enemies. The purpose of this study was to investigate the suitability of crapemyrtle aphids as a food source for the green lacewing *Chrysoperla rufilabris* (Bermeister). Experiments were conducted to test the effects of crapemyrtle cultivar, mature plant height, and source of germplasm on the life history characteristics of *C. rufilabris.* Crapemyrtle aphids were reared on seven cultivars of crapemyrtle, including Apalachee, Natchez, Sioux, Tuscarora, Lipan, Carolina Beauty, and Byers Wonderful White. Aphids from each cultivar were fed exclusively to green lacewing larvae from egg to adult eclosion. Survivorship by cultivar treatment was determined for developmental stages and growth parameters. Cultivar did not affect larval survival but did affect pupal and total survivorship. Length of pupation was extended by aphids feeding on mid mature height cultivars. Dry mass of larvae and adults were also affected accordingly. Males had lower dry mass than females. Lacewings fed on aphids from tall mature height cultivars developed higher dry mass.

**Highlights**

## 2. Introduction

Many studies have indicated plants can influence insect natural enemies at the third trophic level through nutritional, morphological, or chemical attributes (Malcolm 1990, Clark and Messina 1998, Fuentes-Contreras and Niemeyer 1998, Francis et al. 2001, Messina and Sorenson 2001, Giles et al. 2002a, Giles et al. 2002b, Gonzales et al. 2002, Legrand and Barbosa 2003, Kansman et al. 2025) Because tritrophic interactions can have a considerable impact on biological control systems, it is imperative to consider these interactions when implementing or evaluating biological control agents (Price et al. 1980, Van Emden 1995, Bottrell et al. 1998, Vidal and Tscharntke 2001).

One of the best known tritrophic pathways involves aphids and aphid predators, where plants indirectly affect the fitness of aphid predators through the aphid prey (Van Emden 1995). Rothschild et al. (1970) demonstrated that the oleander aphid, *Aphis nerii* Fonscolombe (Hemiptera: Aphididae), can sequester cardenolides from feeding on oleander or milkweed. Further studies indicated that lacewing predators suffered higher mortality rates on *A. nerii* that contained cardenolides and the mortality rate was related to the concentration of cardenolides inside the plant (Rothschild et al. 1970, Malcolm 1990, Hodek 1993, Sohail et al. 2019). Other herbivorous insects protect their offspring from lacewing predation by provisioning their eggs with chemical protections that are derived from their diet. The green lacewing *Ceraeochysa cubana* Hagen, avoids *Utetheisa ornatrix*, (L.) eggs provisioned with alkaloids, but will devour eggs that are left unprotected (Eisner et al. 2000). In addition to allelochemicals, plants affect predators through nutritional pathways. Pea aphids raised on alfalfa and fava bean differ in their myristic acid content, and green lacewing larvae (*Chrysoperla rufilabris* Burmeister) that fed on these aphids exhibited variations in survival rates and adult body mass depending on the aphids’ host plant (Giles et al. 2000).

Green lacewings (Chrysopidae: Neuroptera) have been implicated as an important natural enemy in several biological control systems (New 1975, Liao et al. 1984, Liao et al. 1985, Wang and Nordlund 1994, Randolph et al. 2002, Stewart et al. 2002, Al-Eryan et al. 2024, Karkuzhali et al. 2025). In pecans, *C. rufilabris* is considered to be a key natural enemy and one of the most dominant predators for controlling three species of pecan aphids (Dinkins et al. 1994, Kunkel and Cottrell 2007). Pecan aphids display signs of pesticide resistance, and pesticide applications reduce the number of natural enemies in pecans increasing the need for further pesticide applications to control primary and secondary pests (Mizell and Schiffhauer 1989, Mizell and Sconyers 1992, Mizell 2003). The use of cover crops to increase natural enemy recruitment and retention was suggested in pecans by Tedders (1983), and Mizell and Schiffhauer (1987b). Tedders (1983) suggested using clover and vetch during the spring months when pecan aphid populations are low in order to retain lacewings and other natural enemies that overwinter on the trunks of pecans (Mizell and Schiffhauer 1987a). Mizell and Schiffhauer (1987b) suggested the use of crapemyrtle as an additional augmentation crop because crapemyrtle aphids *Tinocallis kahawaluokalani* (Kirkaldy) have a peak population in the mid to late summer, which is when pecan aphid populations experience a seasonal collapse before increasing again in the fall.

Given this context, the present study aims to determine whether crapemyrtle cultivars influence the suitability of aphid prey for *C. rufilabris.* We propose that aphids feeding on different crapemyrtle cultivars will vary in nutritional or chemical composition, thereby affecting the development, survival, or fitness of their lacewing predators. Understanding how plant cultivar affects this predator-prey dynamic could enhance the effectiveness of conservation biological control strategies in pecan orchards, particularly during seasonal gaps in pest populations. This hypothesis builds on existing evidence of plant-mediated effects on natural enemies (Price et al. 1980; Bottrell et al. 1998; Giles et al. 2002a) and is especially relevant given the proposed role of crapemyrtle as an augmentation crop to sustain beneficial predator populations (Tedders 1983; Mizell and Schiffhauer 1987b).

To evaluate the suitability of crapemyrtle aphids as a food source for the green lacewing *Ceraeochrysa rufilabris*, we examined the effects of crapemyrtle cultivar, mature plant height, and germplasm source on lacewing life history traits. In the first experiment, lacewing larvae were fed *ad libitum* on aphids from seven crapemyrtle cultivars to assess diet quality. In the second experiment, larvae received suboptimal diets consisting of aphids and lepidopteran eggs to determine whether aphids from different cultivars differed in their nutritional or chemical suitability.

## 3. Materials and Methods

### 3.1. Ad libitum Experiment

#### Aphid Rearing

Crapemyrtle aphids (*Tinocallis kahawaluokalani*) were reared on seven crapemyrtle cultivars: ‘Apalachee’, ‘Natchez’, ‘Sioux’, ‘Tuscarora’, ‘Lipan’, ‘Carolina Beauty’, and ‘Byers Wonderful White’. Germplasm sources of these cultivars are described in Herbert et al. (2009). Crapemyrtle cultivars vary in their susceptibility to aphid infestation (Mizell and Knox 1993; Wilson and Bertone 2024). Such variation is relevant to biological control because cultivar-dependent differences in aphid abundance and host suitability may influence prey availability for natural enemies. However, differences in field aphid abundance may also reflect plant characteristics and natural-enemy activity and do not necessarily indicate differences in aphid nutritional quality. Therefore, the present study evaluated the suitability of aphids reared on different crapemyrtle cultivars as prey for the green lacewing *Ceraeochrysa rufilabris* under controlled laboratory conditions.

Aphids were reared on crapemyrtle plants by enclosing individual branches in sleeve cages that were constructed from cloth mesh with 1 mm^2^ openings. Sleeve cages were 16 cm in diameter and 80 cm in length. Branches with sleeve cages had the terminal blooming portion removed before sliding the sleeve over the branch and securing it at both ends using a piece of nylon string that was sewn into the sleeve. To prevent invasion by fire ants (*Solenopsis invicta* Buren), all branches touching the sleeve cage were pruned, and the portion of the branch 10 cm under the sleeve cage was covered with the sticky material Tanglefoot® (Grand Rapids, MI). Sleeve cages were inoculated with aphid-infested leaf disks taken from the laboratory colony being reared on “Tonto” and inoculated two weeks before starting the ad libitum feeding experiment. Four plants per cultivar were used and several branches per plant received sleeve cages to ensure a sufficient supply of aphids.

#### Lacewing Rearing

Green lacewing (*C. rufilabris)* eggs were ordered from Rincon Vitova insectaries (Ventura, CA). Eggs were examined under a microscope to check for signs of development and 400 eggs that appeared healthy were placed singly into cells of 96-well plates. Wells were sealed with Parafilm® that was pressed firmly against the top of the plate to contain first instar larvae and prevent cannibalism. Plates were examined every 24 hr for 1^st^ instar larvae, and upon occurrence, the larvae were placed into a condiment cup that had an organdy cloth-vented lid. As the larvae hatched from their eggs, they were randomly assigned to one of the 28 plants (4 replicates of the 7 cultivars) and given a piece of leaf material infested with aphids. Ten larvae were assigned to each plant (40 per cultivar).

Lacewing larvae were reared in small condiment cups and fed crapemyrtle aphids ad libitum by placing aphid infested leaf material from the appropriate plant directly into each container. During the feeding procedures a small piece of wet cotton was placed into each cup to provide water and maintain a higher humidity than in the growth chamber. The larvae were kept in an environmental growth chamber at 25 ± 1°C with a 16:8 light: dark photoperiod. For each lacewing, the dates of larval emergence, pupation and adult emergence were recorded. On the day of adult emergence, adult lacewings were placed into the freezer at −4 °C. Adults were dried in an oven at 60 °C for 12 h and the dry mass was measured to an accuracy of ± 0.1 mg. The average of three measurements was used as the dry mass in statistical analyses.

The experiment was originally structured as a randomized complete block design, with four aphid-infested plants per cultivar and ten *C. rufilabris* larvae assigned to each plant. However, due to the loss of some sleeve cages caused by branch breakage, weather conditions, farm equipment, or other accidental disturbances, leaf material from additional plants within the same cultivar was used to maintain a consistent food supply for the developing larvae. Although all larvae continued to receive aphids reared on their assigned cultivar, the individual plant could no longer serve as a valid blocking criterion. Consequently, survivorship was analyzed as a binary response variable, with individual lacewings considered as replicates rather than using plants as the unit of replication.

### 3.2. Suboptimal Diets Experiment

#### Aphid rearing

Crapemyrtle aphids were reared in the laboratory on the cultivars, including Byers Wonderful White, Carolina Beauty, Apalachee, and Sioux. Excised leaf tissue, in the form of 40 mm leaf disks, was used to rear crapemyrtle aphids and kept inside 100 x 15 mm water agar Petri dishes. Preparation of agar plates followed the procedures outlined by Herbert et al. (2009); however, 20 ml of agar was used in each 100 x 15 mm Petri plate, instead of 5 ml used in 40 x 10 mm Petri plates. Leaf material was collected in the field by placing leaves into plastic bags and transporting them immediately to the laboratory in a cooler with ice packs. In the laboratory, a 40 mm leaf disk was excised from each leaf and rinsed under cold tap water for several minutes. During rinsing, the disks were gently rubbed between the thumb and forefinger to remove any foreign debris, insects, or mites. Based on results from a previous feeding experiment, where no lacewing larvae survived to adulthood, leaf material used in this study was not disinfected with bleach. Mold was not an issue in this experiment, as the leaf disks were used for only five days. Four leaf disks from the same cultivar were placed in each 100 × 15 mm Petri dish, with the adaxial surface in contact with the agar. Sixty leaf disks per cultivar were prepared each day for use in the experiment.

The crapemyrtle aphid culture was started from adult virginoparae collected from the cultivar “Tonto” in the field. Crapemyrtle branches infested with crapemyrtle aphids were brought into the laboratory and placed in a large container. Sixteen virginoparous aphids, randomly aspirated from the entire pool of field-collected individuals, were added to each Petri dish containing leaf disks. To maintain a culture of nymphs of uniform age and relatively consistent density, the adult aphids were removed every 24 hours and transferred to a new dish containing freshly prepared leaf material from the same cultivar. Aphid nymphs developed into adults within 6-7 days, and the adults obtained after one generation were used to maintain cultivar-specific aphid cultures. Petri dishes containing the aphid cultures were kept on a laboratory bench at 27 ± 2 °C and provided with supplemental lighting. Fluorescent lights were used to maintain a 16:8 light:dark photoperiod. Each fixture contained two 40 W fluorescent tubes and was suspended 0.5 m above the bench surface.

#### Lacewing rearing

Lacewing eggs were obtained from Beneficial Insectary (Redding, CA). Upon arrival, the eggs were inspected, and 132 showing signs of advanced development were individually placed into ¼-dram vials fitted with cotton plugs. Each egg was randomly assigned to one of 11 treatments. Three treatments served as mortality controls, in which lacewing larvae were fed 2.5, 5, or 10 mg of *Ephestia kuehniella* Zeller eggs per day. The remaining eight treatments tested the effects of prey level and cultivar. In these treatments, larvae were fed either 25 or 50 aphids per day reared on one of four crapemyrtle cultivars: Apalachee, Sioux, Carolina Beauty, or Byers Wonderful White.

During feeding, larvae were removed from the vials, and the vials were cleaned using a pipe cleaner to remove excrement and any remaining prey. If a larva was attached to the vial and in the process of molting, it was not removed. Instead, remaining prey were carefully extracted using a small camel-hair brush, avoiding any contact with the larva. Most larvae molted on the cotton plug, which allowed them to be removed without disturbance.

Aphids were delivered to the vials by first aspirating them into a 1.5 mL Eppendorf tube. The vial was then inserted into the tube until a seal was formed, and the aphids were gently tapped into the vial. This method of transfer was found to be non-lethal to the aphids and more efficient than using a brush. *Ephestia* eggs were measured and delivered volumetrically by calibrating a 10 µL pipette tip to correspond to the three diet levels. The 10 mg *Ephestia* egg treatment was considered an ad libitum diet, as uneaten eggs were consistently present throughout all larval instars.

Vials were placed on their sides in the lid of a large 150 × 20 mm Petri plate. To maintain a relative humidity of 60-80%, two pieces of cotton moistened with deionized water were placed inside the plate, and the base of the Petri dish was set slightly ajar on the lid. Humidity was monitored periodically in each Petri plate and found to consistently remain within the target range.

### 3.3. Experimental Parameters

Two samples of 25 aphids were collected from each cultivar and placed into pre-weighed 1.5 mL Eppendorf tubes. The fresh mass of each sample was determined by weighing the tube again after the aphids were added. Following weighing, the samples were stored in a freezer at −4 °C. For each lacewing, the dates of larval eclosion, pupation, and adult emergence were recorded. Pupal mass was measured three days after pupation began, and adult fresh mass was recorded on the day of emergence. After recording fresh mass, adult lacewings were stored at −4 °C. Adult lacewings and aphid samples were lyophilized by first freezing them at −80 °C for two hours, then immediately transferring them to a Labconco FreeZone 2.5 lyophilizer for 24 hours. The dry mass of aphid and lacewing samples was measured to an accuracy of ±0.01 mg.

### 3.4. Statistical Analyses

Analyses were conducted in SAS using PROC GLIMMIX (SAS Institute, 2000–2004). Survivorship data were analyzed assuming a binomial distribution with the default logit link function. Larval development time and adult dry mass were analyzed using a lognormal distribution. Pupal development time was analyzed after applying a square root transformation, while pupal mass data did not require transformation.

## 4. Results

### 4.1. Ad libitum Experiment

The cultivar or source of aphid prey significantly affected pupal survivorship, and total survivorship (Table 1; Fig. 1A-B). Pupal survivorship was lower for lacewings fed aphids reared on “Apalachee” compared to those fed aphids from other *L. fauriei* germplasm sources (Table 1; Fig. 1A). Within the “Basham’s Party Pink” *L. fauriei* germplasm source, pupal emergence was lower in lacewings fed aphids reared on “Lipan,” a cultivar characterized by medium mature plant height (Table 1; Fig. 1A).

**Figure 1.**
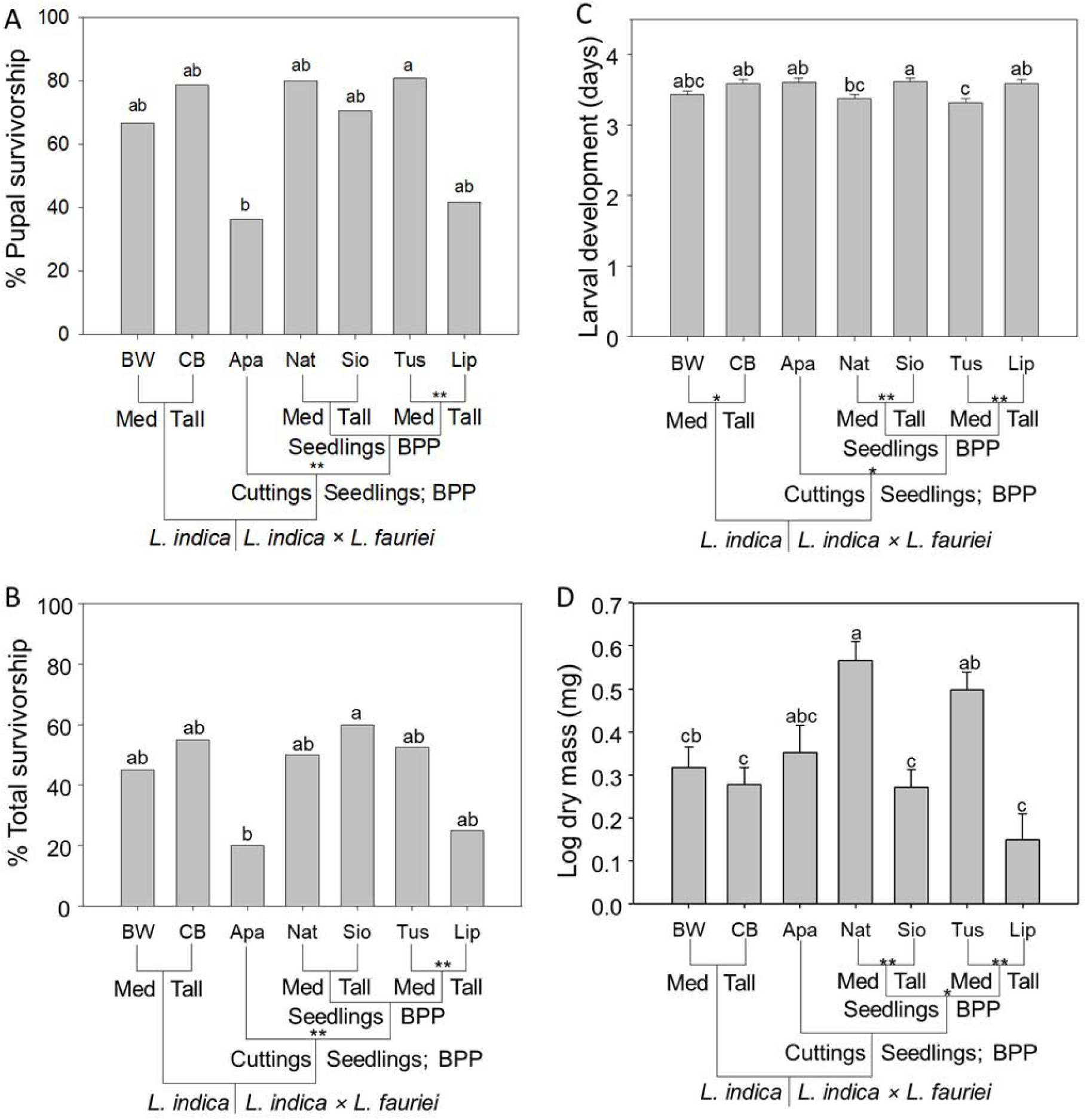
Percentage of lacewing pupae that emerged as adults (A), percentage of lacewing larvae that successfully developed from larva to adult (B), least square mean (± SE) of the square root–transformed larval development time in days (C), and least square mean (± SE) of the log–transformed dry mass in mg for adult lacewings (D) in the ad libitum diet experiment. Data are shown for the cultivars ‘Byers Wonderful White’ (BW), ‘Carolina Beauty’ (CB), ‘Apalachee’ (Apa), ‘Natchez’ (Nat), ‘Sioux’ (Sio), ‘Tuscarora’ (Tus), and ‘Lipan’ (Lip). Bars labeled with different letters differ significantly according to Tukey’s HSD (*P* < 0.05). Orthogonal contrasts are indicated by asterisks: *P* < 0.05 (*), P < 0.01 (**).

**Table 1.**
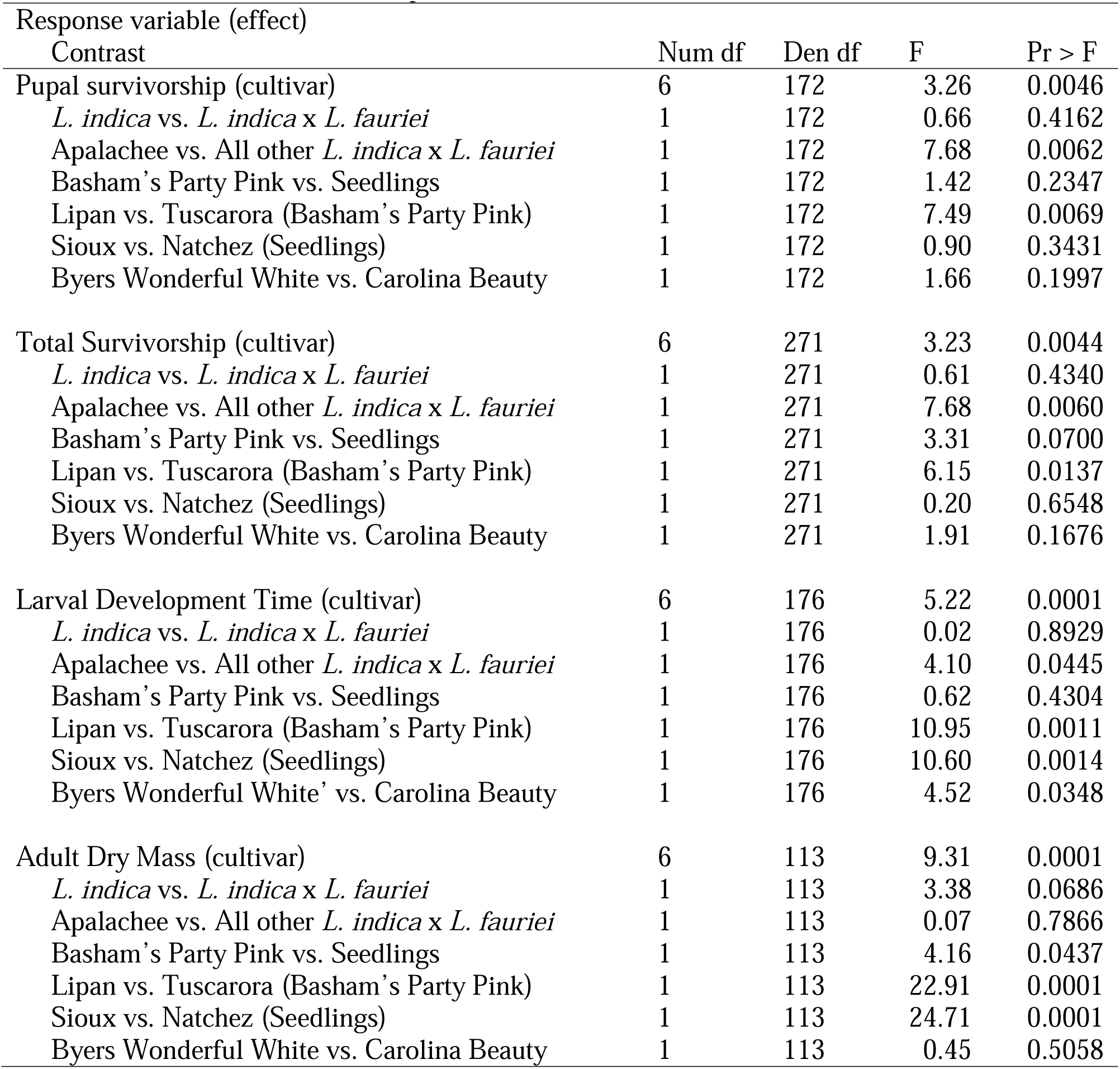
Analysis of variance results for the ad libitum diet experiment with significant variables and contrast statements performed in each test.

| Response variable (effect) |  |  |  |  |
| Contrast | Num df | Den df | F | Pr > F |
|---|---|---|---|---|
| Pupal survivorship (cultivar) | 6 | 172 | 3.26 | 0.0046 |
| *L. indica* vs. *L. indica* x *L. fauriei* | 1 | 172 | 0.66 | 0.4162 |
| Apalachee vs. All other *L. indica* x *L. fauriei* | 1 | 172 | 7.68 | 0.0062 |
| Basham’s Party Pink vs. Seedlings | 1 | 172 | 1.42 | 0.2347 |
| Lipan vs. Tuscarora (Basham’s Party Pink) | 1 | 172 | 7.49 | 0.0069 |
| Sioux vs. Natchez (Seedlings) | 1 | 172 | 0.90 | 0.3431 |
| Byers Wonderful White vs. Carolina Beauty | 1 | 172 | 1.66 | 0.1997 |
| Total Survivorship (cultivar) | 6 | 271 | 3.23 | 0.0044 |
| *L. indica* vs. *L. indica* x *L. fauriei* | 1 | 271 | 0.61 | 0.4340 |
| Apalachee vs. All other *L. indica* x *L. fauriei* | 1 | 271 | 7.68 | 0.0060 |
| Basham’s Party Pink vs. Seedlings | 1 | 271 | 3.31 | 0.0700 |
| Lipan vs. Tuscarora (Basham’s Party Pink) | 1 | 271 | 6.15 | 0.0137 |
| Sioux vs. Natchez (Seedlings) | 1 | 271 | 0.20 | 0.6548 |
| Byers Wonderful White vs. Carolina Beauty | 1 | 271 | 1.91 | 0.1676 |
| Larval Development Time (cultivar) | 6 | 176 | 5.22 | 0.0001 |
| *L. indica* vs. *L. indica* x *L. fauriei* | 1 | 176 | 0.02 | 0.8929 |
| Apalachee vs. All other *L. indica* x *L. fauriei* | 1 | 176 | 4.10 | 0.0445 |
| Basham’s Party Pink vs. Seedlings | 1 | 176 | 0.62 | 0.4304 |
| Lipan vs. Tuscarora (Basham’s Party Pink) | 1 | 176 | 10.95 | 0.0011 |
| Sioux vs. Natchez (Seedlings) | 1 | 176 | 10.60 | 0.0014 |
| Byers Wonderful White’ vs. Carolina Beauty | 1 | 176 | 4.52 | 0.0348 |
| Adult Dry Mass (cultivar) | 6 | 113 | 9.31 | 0.0001 |
| *L. indica* vs. *L. indica* x *L. fauriei* | 1 | 113 | 3.38 | 0.0686 |
| Apalachee vs. All other *L. indica* x *L. fauriei* | 1 | 113 | 0.07 | 0.7866 |
| Basham’s Party Pink vs. Seedlings | 1 | 113 | 4.16 | 0.0437 |
| Lipan vs. Tuscarora (Basham’s Party Pink) | 1 | 113 | 22.91 | 0.0001 |
| Sioux vs. Natchez (Seedlings) | 1 | 113 | 24.71 | 0.0001 |
| Byers Wonderful White vs. Carolina Beauty | 1 | 113 | 0.45 | 0.5058 |

Total survivorship from larva to adult followed the same pattern as pupal survivorship, showing significant differences associated with the source of *L. fauriei* germplasm (Table 1; Fig.1B) and between the cultivars “Lipan” and “Tuscarora” (Table 1; Fig.1B).

Lacewing larvae fed aphids reared on “Apalachee” required a longer time to reach pupation compared to those fed aphids from other *L. fauriei* germplasm sources (Table 1; Fig. 1C). Within the pure *L. indica*, *L. fauriei* seedlings, and *L. fauriei* “Basham’s Party Pink” germplasm sources, cultivar mature plant height influenced larval development. In each source, larvae fed aphids reared on cultivars with medium mature plant height took longer to pupate (Table 1; Fig. 1C).

Adult dry mass was greater for lacewings fed aphids from cultivars in the *L. fauriei* seedlings germplasm source compared to those fed aphids from the *L. fauriei* “Basham’s Party Pink” source (Table 1; Fig. 1D). Additionally, adult dry mass differences were associated with mature plant height, with lacewings fed aphids from cultivars of tall mature plant height exhibiting higher dry mass than those fed aphids from cultivars with medium mature plant height (Table 1; Fig. 1D)

### 4.2. Suboptimal Diet Experiment

#### Aphid diets

Aphids reared on pure *L. indica* cultivars had a lower dry mass than aphids reared on the *L. indica* × *L. fauriei* cultivars (Table 2; Fig. 2A). Aphid dry mass was linearly related to fresh mass for all cultivars (Table 3; Fig. 2B). Slopes for the regressions of dry mass vs. fresh mass were not parallel, leading to an analysis that compared the ratio of dry mass to fresh mass among the four cultivars tested. The ratio of dry mass to fresh mass for aphids was different between cultivars “Byers Wonderful White” and “Sioux” (F=3.75; df=3, 40; P=0.0182).

**Figure 2.**
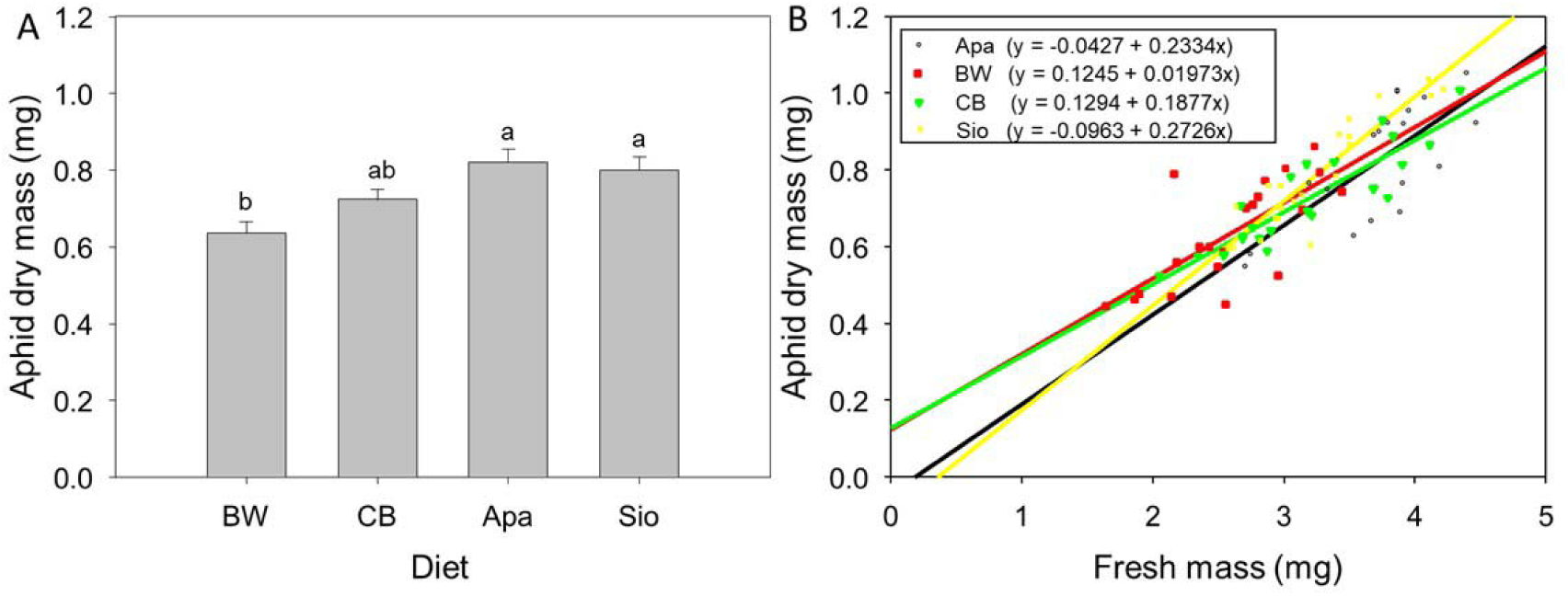
(A) Mean dry mass (± SE) of crapemyrtle aphid samples in the suboptimal diet experiment for the cultivars ‘Byers Wonderful White’ (BW), ‘Carolina Beauty’ (CB), ‘Apalachee’ (Apa), and ‘Sioux’ (Sio). Bars labeled with different letters are significantly different according to Tukey’s HSD (*P* < 0.05). (B) Scatter plots with regression lines showing the relationship between dry mass and fresh mass of aphid samples collected from the same cultivars.

**Table 2.** Analysis of variance results with contrasts for the dry mass of crapemyrtle aphids reared on different cultivars of crapemyrtle.

| Response Contrast | Num df | Den df | F | Pr > F |
|---|---|---|---|---|
| Dry Mass (cultivar) | 3 | 40 | 11.42 | <0.0001 |
| *L. indica* vs. *L. indica* x *L. fauriei* | 1 | 40 | 20.74 | <0.0001 |
| Medium vs. Tall | 1 | 40 | 0.43 | 0.5175 |

**Table 3.**
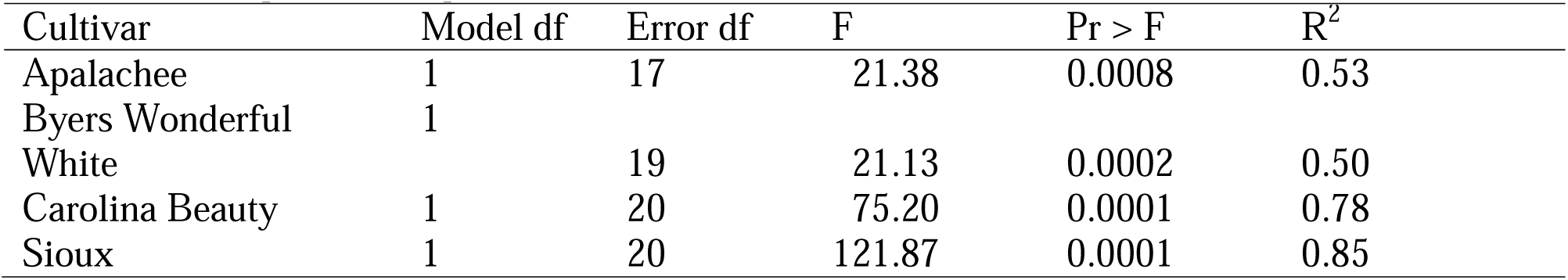
Regression statistics for the comparison of aphid dry mass to aphid fresh mass using samples of 25 aphids.

| Cultivar | Model df | Error df | F | Pr > F | R<sup>2</sup> |
|---|---|---|---|---|---|
| Apalachee | 1 | 17 | 21.38 | 0.0008 | 0.53 |
| Byers Wonderful White | 1 | 19 | 21.13 | 0.0002 | 0.50 |
| Carolina Beauty | 1 | 20 | 75.20 | 0.0001 | 0.78 |
| Sioux | 1 | 20 | 121.87 | 0.0001 | 0.85 |

#### Lacewing diets

Larval developmental time was shorter for lacewing larvae that were given higher levels of daily prey, and this effect was consistent for all crapemyrtle cultivars and the *Ephestia* egg controls (Table 4). In addition to differences in daily prey levels, larvae reared on *Ephestia* eggs had a lower duration of larval development in comparison to larvae reared on crapemyrtle aphids (Fig. 3A). Pupal duration was shorter for larvae fed *Ephestia* eggs, however, duration did not differ for lacewing larvae fed aphids from different crapemyrtle cultivars (Table 4; Fig. 3B).

**Figure 3.**
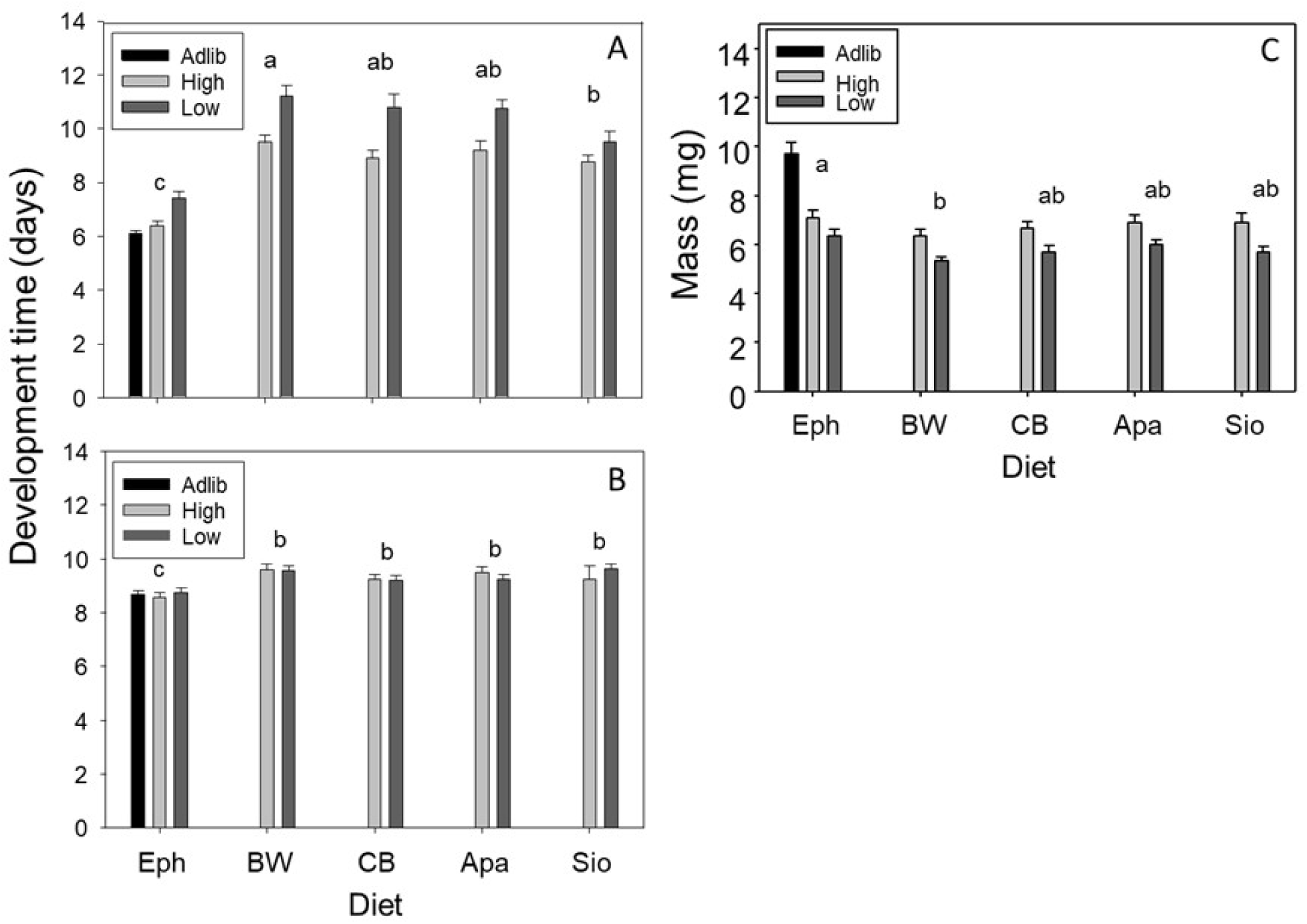
Mean developmental time (± SE) in days for the suboptimal diet experiment for (A) lacewing larvae and (B) lacewing pupae, and (C) mean pupal mass (± SE) of lacewings reared on different diet sources: *Ephestia* eggs (Eph) and crapemyrtle aphids from the cultivars ‘Byers Wonderful White’ (BW), ‘Carolina Beauty’ (CB), ‘Apalachee’ (Apa), and ‘Sioux’ (Sio). Paired columns (low and high prey levels) sharing the same letter are not significantly different; bars with different letters indicate significant differences among diets based on Tukey’s HSD (*P* < 0.05). The Eph *ad* libitum treatment was not included in statistical analyses but is shown for reference to illustrate relative trends.

**Table 4.** Analysis of variance results for *C. rufilabris* in the suboptimal diet experiment. Response.

| Response |  |  |  |  |
|---|---|---|---|---|
| Effect | Num df | Den df | F | Pr > F |
| Larval Development |  |  |  |  |
| Diet Source | 4 | 86 | 46.69 | <0.0001 |
| Prey Level | 1 | 86 | 44.93 | <0.0001 |
| Diet Source × Prey Level | 4 | 86 | 0.70 | 0.5962 |
| Pupal Duration |  |  |  |  |
| Diet Source | 4 | 71 | 7.31 | <0.0001 |
| Prey Level | 1 | 71 | 0.19 | 0.6640 |
| Diet Source × Prey Level | 4 | 71 | 0.67 | 0.6171 |
| Pupal Mass |  |  |  |  |
| Diet Source | 4 | 86 | 2.92 | 0.0257 |
| Prey Level | 1 | 86 | 30.36 | <0.0001 |
| Diet Source × Prey Level | 4 | 86 | 0.21 | 0.9341 |
| Adult Dry Mass |  |  |  |  |
| Diet Source | 4 | 61 | 6.24 | 0.0003 |
| Prey Level | 1 | 61 | 61.34 | <0.0001 |
| Sex | 1 | 61 | 23.41 | <0.0001 |
| Diet Source × Prey Level | 4 | 61 | 1.17 | 0.3324 |
| Diet Source × Sex | 4 | 61 | 1.28 | 0.2872 |
| Sex × Prey Level | 1 | 61 | 0.13 | 0.7189 |

Pupal mass increased with daily prey availability, and this trend was consistent across all crapemyrtle cultivars and the Ephestia egg control (Table 4; Fig. 3C). Lacewing larvae fed Ephestia eggs produced heavier pupae than those fed crapemyrtle aphids from ‘Byers Wonderful White’ (Table 4; Fig. 3C).

Lacewing adult dry mass was different among the crapemyrtle and *Ephestia* egg treatments (Table 4; Fig. 4A). Lacewings provisioned with higher levels of daily prey had a greater dry mass (Table 4; Fig. 4B), and males had a lower dry mass than females (Table 4; Fig. 4C).

**Figure 4.**
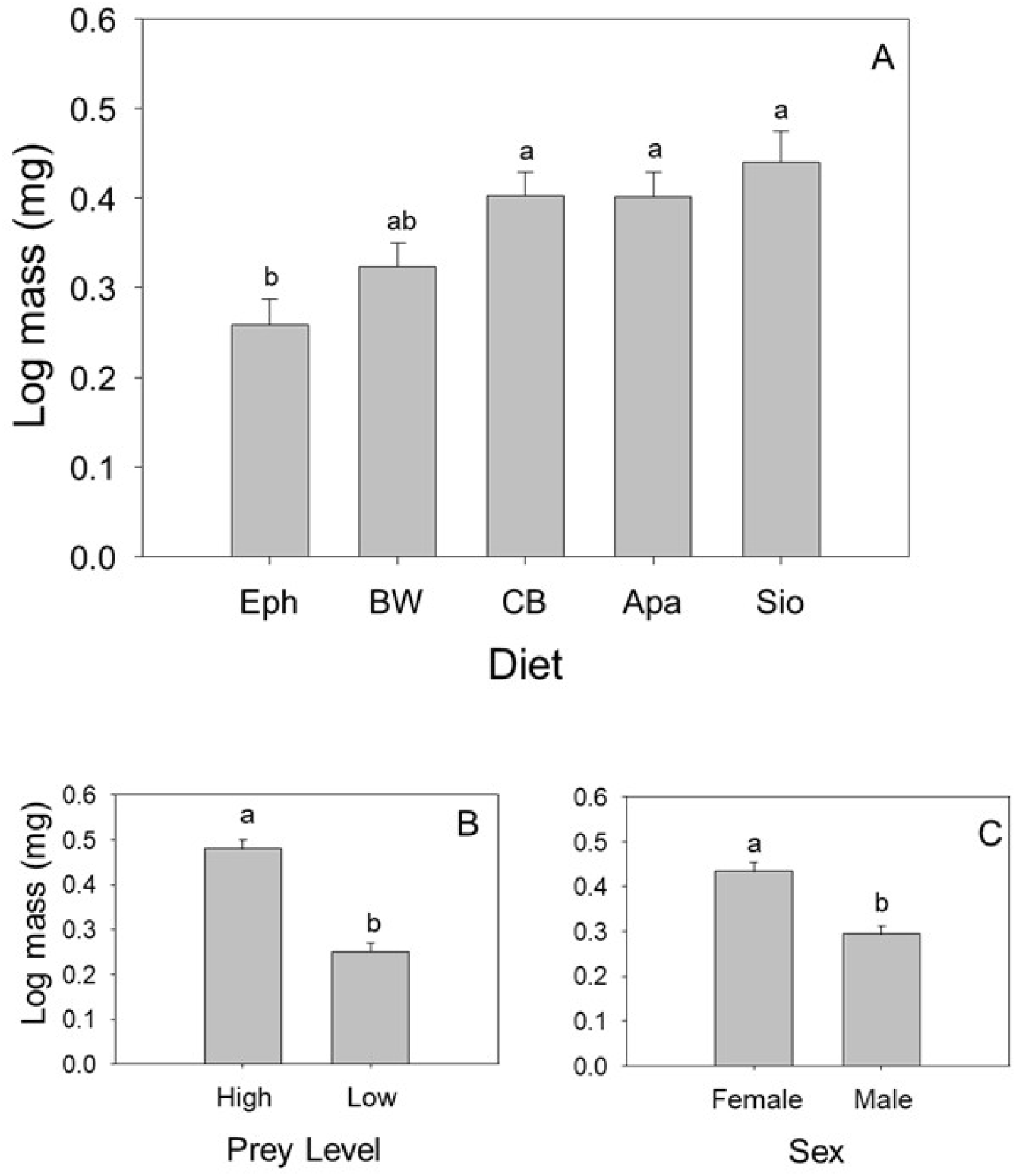
Least square mean (log-transformed) adult dry mass (± SE) of lacewings that successfully emerged without deformation in the suboptimal diet experiment, shown for the main effects of (A) diet, (B) daily prey level, and (C) sex. Diet sources included *Ephestia* eggs (Eph) and crapemyrtle aphids from the cultivars ‘Byers Wonderful White’ (BW), ‘Carolina Beauty’ (CB), ‘Apalachee’ (Apa), and ‘Sioux’ (Sio). Bars labeled with different letters are significantly different based on Tukey’s HSD (*P* < 0.05).

## 5. Discussion

Results from Ad libitum food consumption experiment indicate that there are multitrophic interactions among crapemyrtle, crapemyrtle aphids, and *C. rufilabris*. The pupal stage appears to be the most vulnerable stage with respect to survivorship, and the most critical stage with regards to developmental time was the larval stage. Orthogonal contrasts indicated that lacewings were affected by broader plant attributes particularly mature plant height, and source of *L. fauriei* germplasm. If crapemyrtle is evaluated for enhancing biological control in pecans or other economically important crops as suggested by Mizell and Schiffhauer (1987b), this research may be helpful for selecting the most appropriate cultivars for use as augmentation crops. ‘Natchez’ and ‘Tuscarora’ are tall cultivars from the *L. indica* × *L. fauriei* parentage, and lacewings fed crapemyrtle aphids reared on these cultivars had the shortest development time, lowest pupal mortality, and highest adult dry mass. Previous studies (Herbert et al. 2009) showed that cultivars from the *L. indica* × *L. fauriei* parentage were a more suitable host and more preferred by crapemyrtle aphids. ‘Natchez’ was rarely infested with high populations of crapemyrtle aphids in the field and had the lowest susceptibility rating among the cultivars tested (Mizell and Knox 1993). Lower aphid populations in the field could be due to higher predation rates by lacewings or other aphid predators. This may be a result of either qualitative or quantitative differences in aphid prey.

Our findings demonstrate that *Lagerstroemia* cultivar selection directly shapes the efficacy of *C. rufilabris* as a biological control agent through plant-mediated effects on prey suitability. Critically, cultivars derived from *L. fauriei* germplasm (notably‘Natchez’ and‘Tuscarora’) enhanced lacewing fitness by reducing larval development time, increasing pupal survivorship, and yielding larger adults - traits directly linked to improved predator field performance. This cultivar-dependent variation in prey quality likely stems from underlying differences in nutritional and/or allelochemical properties transmitted through the aphid, as evidenced by differential lacewing responses in both ad libitum and suboptimal feeding contexts. These results validate the strategic use of crapemyrtle as an augmentation crop in pecan systems: selecting high-performing cultivars can sustain robust *C. rufilabris* populations during seasonal gaps in pecan aphid availability, thereby strengthening year-round conservation biological control. The paradoxical suitability of‘Natchez’, a cultivar exhibiting low aphid susceptibility yet high lacewing quality, further highlights its unique value in promoting predator retention without encouraging pest outbreaks.

Qualitative or quantitative differences in aphid prey have been determined by other researchers with suboptimal diets (Giles et al. 2000, Giles et al. 2002a, Giles et al. 2002b). The previous authors found that these differences were due to a specific fatty acid, but predators can be affected by toxic allelochemicals present within the aphids’ host plant (Price et al. 1980, Malcolm 1990, Van Emden 1995). *Lagerstroemia indica* has several alkaloids that are found in the leaves, bark, and seeds (Ferris et al. 1971c, Ferris et al. 1971b, a). Mizell et al. (2002) suggested that these chemicals could play a defensive role and help protect crapemyrtle aphids against insect parasitism. The fact that no parasitoids have been discovered from areas searched that have both the hosts and the crapemyrtle aphid such as Southeast Asia and Latin America also supports this hypothesis (Mizell et al. 2002). Moreover, the difference in aphid numbers on cultivars with different genetic backgrounds and their differential impacts on native predators also suggests that the aphid over evolutionary time has not totally countered the negative effect(s) of plant chemistry. Further experiments examining more homogenous aphid populations at differential diet levels may help in determining what factors are affecting lacewing survival and development, and if these factors are qualitative or quantitative in nature.

In the suboptimal diet experiment, lacewing survivorship was not found to be significantly affected by cultivar or the amount of food provisioned daily. However, daily prey levels and food source were factors in determining larval developmental time, pupal mass and adult dry mass. Lacewing larvae developed faster and had a higher pupal mass when they were reared on higher daily prey levels regardless of the food source. Studies on the diet and life history of *C. rufilabris* have indicated that lower larval developmental time is associated with higher quality food (Hydorn and Whitcomb 1979, Legaspi et al. 1994, Legaspi et al. 1996). However, Hydorn and Whitcomb (1979) also showed that *C. rufilabris* larvae reared on *Tribolium castaneum* (Herbst) had a higher adult mass, but lower fecundity than *C. rufilabris* larvae that were reared on aphids or *Phthorimaea operculella* Zeller eggs. The previous authors attributed the higher adult mass to a longer developmental time, and in this experiment, *C. rufilabris* that were reared on crapemyrtle aphids had a longer larval development time and a larger dry mass than lacewings reared on *Ephestia* eggs. These results are interesting because lacewing larvae reared on *Ephestia* eggs had a higher pupal mass than larvae reared on crapemyrtle aphids and the results of adult dry mass are not consistent with the relationship between pupal mass and adult dry mass. The discrepancy between these findings may be due to pupal survivorship, and although the relationship is not significant, pupae that did not successfully emerge as adults had a marginally higher pupal mass than pupae that did emerge. This relationship can be seen in the survivorship data for lacewings that were fed crapemyrtle aphids, and while the survivorship data is not statistically significant, which may be due to issues with replication, lacewing larvae reared on crapemyrtle aphids from the cultivars ‘Carolina Beauty’, ‘Tuscarora’, and ‘Sioux’ suffered higher mortality when they were fed at the higher daily prey level as perhaps one would observe from toxicant doses.

This study provides the first empirical validation that crapemyrtle cultivar selection is a critical operational variable for enhancing *C. rufilabris* conservation in pecan systems. By demonstrating that plant genetic identity directly modulates lacewing fitness via prey quality, we establish a functional basis for using augmentation crops with precision moving beyond simple presence/absence to strategic cultivar deployment. Future work should prioritize identifying the specific phytochemical or nutritional mechanisms underlying these cultivar effects, enabling the design of multifunctional agroecosystems that concurrently support natural enemies and suppress pest resurgence.

## Author Contributions

John J. Herbert: data curation, investigation, formal analysis, visualization, writing – original draft. Romain Exilien: Writing – review and editing. Heather McAuslane: conceptualization, investigation, funding acquisition, writing – review and editing, validation, methodology, project administration, data curation, supervision, resources. Rusell. F. Mizell, III: conceptualization, investigation, funding acquisition, validation, methodology, writing – review and editing, project administration, data curation, supervision, resources.

## Declaration of Competing Interest

The authors declare that they have no competing interests.

## Acknowledgment

We acknowledge the support of the University of Florida Institute of Food and Agricultural Sciences (UF/IFAS) in the implementation of this project.

## Funding

This research was partially funded by internal support from the University of Florida/IFAS.

## Notes

### Competing Interest Statement

The authors have declared no competing interest.

## References Cited

1. Al-Eryan, M. A., Abu-shall, A. M., Tawfeek, M. E., Ibrahim, S. A. s. 2024. Effect of feeding type on some biological aspects of aphid lion, Chrysoperla carnea (Neuroptera:Chrysopidae) under laboratory conditions. Alexandria Science Exchange Journal. 45(3):21–34. 10.21608/asejaiqjsae.2024.388850

2. Bottrell, D. G., P. Barbosa, and F. Gould. 1998. Manipulating natural enemies by plant variety selection and modification: a realistic strategy. Annu. Rev. Entomol. 43: 347–367.

3. Clark, T. L., and F. J. Messina. 1998. Foraging behavior of lacewing larvae (Neuroptera: Chrysopidae) on plants with divergent architectures. J. Insect Behav. 11: 303–317.

4. Dinkins, R. L., W. L. Tedders, and W. Reid. 1994. Predaceous neuropterans in Georgia and Kansas pecan trees. J. Entomol. Sci. 29: 165–175.

5. Eisner, T., M. Eisner, C. Rossini, V. K. Iyengar, B. L. Roach, E. Benedikt, and J. Meinwald. 2000. Chemical defense against predation in an insect egg. Proc. Natl. Acad. Sci. U.S.A.97: 1634–1639.

6. Ferris, J. P., R. C. Briner, and C. B. Boyce. 1971a. Lythraceae alkaloids. VIII. the structure and stereochemistry of the biphenyl ether alkaloids from Decodon *verticillatus* J. Am. Chem. Soc. 93: 2953–2957.

7. Ferris, J. P., R. C. Briner, and C. B. Boyce. 1971b. Lythraceae alkaloids. IX. the isolation and structure elucidation of the alkaloids of Lagerstroemia *indica* L. J. Am. Chem. Soc. 93: 2858–2962.

8. Ferris, J. P., C. B. Boyce, R. C. Briner, U. Weiss, I. H. Qureshi, and N. E. Sharpless. 1971c. Lythraceae alkaloids. X. assignment of absolute stereochemistries on the basis of chiraloptical effects. J. Am. Chem. Soc. 93: 2963–2968.

9. Francis, F., E. Haubruge, P. Hastir, and C. Gaspar. 2001. Effect of aphid host plant on development and reproduction of the third trophic level, the predator *Adalia bipunctata* (Coleoptera: Coccinellidae). Environ. Entomol. 30: 947–952.

10. Fuentes-Contreras, E., and H. M. Niemeyer. 1998. Dimboa glucoside, a wheat chemical defense, affects host acceptance and suitability of *Sitobion avenae* to the cereal aphid parasitoid *Aphidius rhopalosiphi*. J. Chem. Ecol. 24: 371–381.

11. Giles, K. L., R. D. Madden, M. E. Payton, and J. W. Dillwith. 2000. Survival and development of Chrysoperla *rufilabris* (Neuroptera: Chrysopidae) supplied with pea aphids (Homoptera: Aphididae) reared on alfalfa and fava bean. Environ. Entomol. 29: 304–311.

12. Giles, K. L., R. C. Berberet, A. A. Zarrabi, and J. W. Dillwith. 2002a. Influence of alfalfa cultivar on suitability of Acyrthosiphon *kondoi* (Homoptera: Aphididae) for survival and development of *Hippodamia convergens* and Coccinella *septempunctata* (Coleoptera: Coccinellidae). J. Econ. Entomol. 95: 552–557.

13. Giles, K. L., R. D. Madden, R. Stockland, M. E. Payton, and J. W. Dillwith. 2002b. Host plants affect predator fitness via the nutritional value of herbivore prey: investigation of a plant-aphid-ladybeetle system. BioControl (Dordrecht) 47: 1–21.

14. Gonzales, W. L., E. Fuentes-Contreras, and H. M. Niemeyer. 2002. Host plant and natural enemy impact on cereal aphid competition in a seasonal environment. Oikos 96: 481–491.

15. Hodek, I. 1993. Habitat and food specificity in aphidophagous predators. Biocontrol Sci. Technol. 3: 91–100.

16. Hydorn, S. B., and W. H. Whitcomb. 1979. Effects of larval diet on *Chrysopa rufilabris*. Fla. Entomol. 62: 293–298.

17. Kansman, J. T., Hermann, S. L., Ali, J. G., et al. 2025. Flipping indirect defense: chemical cues from natural enemies mediate multitrophic interactions. Current Opinion in Insect Science. 68:101330. 10.1016/j.cois.2024.101330

18. Karkuzhali N, Edward YSJT, Chitra N, et al. 2025. Unveiling the diversity of gut microbes in green lacewings (Chrysopidae: Neuroptera) and their role as protagonist in nutrition. Arch Microbiol. 207(5):100. 10.1007/s00203-025-04289-0

19. Kunkel BA, Cottrell TE. 2007. Oviposition Response of Green Lacewings (Neuroptera: Chrysopidae) to Aphids (Hemiptera: Aphididae) and Potential Attractants on Pecan. Environ. Entomol. 36(3):577–583. https://doi.org/10.1603/0046-225X(2007)36[577:OROGLN]2.0.CO;2

20. Legaspi, J. C., R. I. Carruthers, and D. A. Nordlund. 1994. Life history of *Chrysoperla rufilabris* (Neuroptera: Chrysopidae) provided sweetpotato whitefly *Bemisia tabaci* (Homoptera: Aleyrodidae) and other food. Biol. Control 4: 178–184.

21. Legaspi, J. C., D. A. Nordlund, and B. C. Legaspi, Jr. 1996. Tri-trophic interactions and predation rates in *Chrysoperla* spp. attacking the silverleaf whitefly. Southwest. Entomol. 21: 33–42.

22. Legrand, A., and P. Barbosa. 2003. Plant morphological complexity impacts foraging efficiency of adult *Coccinella septempunctata* L. (Coleoptera: Coccinellidae). Environ. Entomol. 32: 1219–1226.

23. Liao, H. T., M. K. Harris, F. E. Gilstrap, and F. Mansour. 1985. Impact of natural enemies on the blackmargined pecan aphid, *Monellia caryella* (Homoptera: Aphidae). Environ. Entomol. 14: 122–126.

24. Liao, H. T., M. K. Harris, F. E. Gilstrap, D. A. Dean, C. W. Agnew, G. J. Michels, and F. Mansour. 1984. Natural enemies and other factors affecting seasonal abundance of the blackmargined aphid on pecan. Southwest. Entomol. 9: 404–420.

25. Malcolm, S. B. 1990. Chemical defence in chewing and sucking insect herbivores: plant-derived cardenolides in the monarch butterfly and oleander aphid. Chemoecology 1: 12–21.

26. Messina, F. J., and S. M. Sorenson. 2001. Effectiveness of lacewing larvae in reducing Russian wheat aphid populations on susceptible and resistant wheat. Biol. Control 21: 19–26.

27. Mizell, R. F., III. 2003. Summary and overview of pecan arthropod pest management. Southwest. Entomol. 27: 135–140.

28. Mizell, R. F., III, and D. E. Schiffhauer. 1987a. Trunk traps and overwintering predators in pecan orchards - survey of species and emergence times. Fla. Entomol. 70: 238–244.

29. Mizell, R. F., III, and D. E. Schiffhauer. 1987b. Seasonal abundance of the crapemyrtle aphid, *Sarucallis kahawaluokalani*, in relation to the pecan aphids, *Monellia caryella* and *Monelliopsis pecanis* and their common predators. Entomophaga 32: 511–520.

30. Mizell, R. F., III, and D. E. Schiffhauer. 1989. Effects of pesticides on pecan aphid predators *Chrysoperla rufilabris* (Neuroptera: Chrysopidae), *Hippodamia convergens, Cycloneda sanguinea* (L.), *Olla v-nigrum* (Coleoptera: Coccinellidae), and *Aphelinus perpallidus* (Hymenoptera: Encyrtidae). J. Econ. Entomol. 83: 1806–1812.

31. Mizell, R. F., III, and M. C. Sconyers. 1992. Toxicity of imidacloprid to selected arthropod predators in the laboratory. Fla. Entomol. 75: 277–280.

32. Mizell, R. F., III, and G. W. Knox. 1993. Susceptibility of crapemyrtle, *Lagerstroemia indica* L., to the crapemyrtle aphid (Homoptera: Aphididae) in North Florida. J. Entomol. Sci. 28: 1–7.

33. Mizell, R. F., III, F. D. Bennett, and D. K. Reed. 2002. Unsuccessful search for parasites of the crapemyrtle aphid, *Tinocallis kahawaluokalani* (Homoptera: Aphididae). Fla. Entomol. 85: 521–523.

34. New, T. R. 1975. The biology of Chrysopidae and Hemerobiidae (Neuroptera), with reference to their usage as biological control agents: a review. Trans. R. Ent. Soc. Lond. 127: 115–140.

35. Price, P. W., C. E. Bouton, P. Gross, B. A. McPheron, J. N. Thompson, and A. E. Weis. 1980. Interactions among three trophic levels: influence of plants on interactions between insect herbivores and natural enemies. Annu. Rev. Ecol. Syst. 11: 41–65.

36. Randolph, T. L., M. K. Kroening, J. B. Rudolph, F. B. Peairs, and R. F. Jepson. 2002. Augmentative releases of commercial biological control agents for Russian wheat aphid management in winter wheat. Southwest. Entomol. 27: 37–44.

37. Rothschild, M., J. Von Euw, and T. Reichstein. 1970. Cardiac glycosides in the oleander aphid *Aphis nerii*. J. Insect Physiol. 16: 1141–1145.

38. SAS. Institute Inc., SAS 9.1.3 Help and Documentation, Cary, NC: SAS Institute Inc., 2000-2004.

39. Sohail M, Muhammad R, Soomro Q. 2019. Effect of oleander aphid (*Aphis nerii* Boyer de Fonscolombe) on the Mortality and Biological Parameters of Green Lacewing (*Chrysoperla carnea* Stephen). Pakistan Journal of Zoology. 52(1):231–237. 10.17582/journal.pjz/2020.52.1.231.237

40. Stewart, C. D., S. K. Braman, and A. F. Pendley. 2002. Functional response of the azalea plant bug (Heteroptera: Miridae) and a green lacewing *Chrysoperla rufilabris* (Neuroptera: Chrysopidae), two predators of the azalea lace bug (Heteroptera: Tingidae). Environ. Entomol. 31: 1184–1190.

41. Tedders, W. L. 1983. Insect management in deciduous orchard ecosystems: habitat manipulation. Environ. Manage. 7: 29–34.

42. Van Emden, H. F. 1995. Host plant-Aphidophaga interactions. Agriculture Ecosystems and Environment 52: 3–11.

43. Vidal, S., and T. Tscharntke. 2001. Multitrophic plant-insect interactions. Basic Appl. Ecol. 1: 1–2.

44. Wang, R., and D. A. Nordlund. 1994. Use of *Chrysoperla* spp. (Neuroptera: Chrysopidae) in augmentative release programs for control of arthropod pests. Biocontrol News and Information 15: 51N–57N.

45. Wilson CJ, Bertone MA. 2024. Ecology and management of the crapemyrtle aphid (Hemiptera: Aphididae) on crapemyrtle (Myrtales: Lythraceae) in the southern United States. Journal of Integrated Pest Management. 15(1):11. 10.1093/jipm/pmae003

